# Structure of a consensus chitin-binding domain revealed by solution NMR

**DOI:** 10.1101/2020.01.08.899344

**Authors:** Dario Heymann, Harini Mohanram, Akshita Kumar, Chandra S. Verma, Julien Lescar, Ali Miserez

**Affiliations:** Biological and Biomimetic Material Laboratory, Center for Biomimetic Sensor Science, School of Materials Science and Engineering, Nanyang Technological University (NTU), 50 Nanyang Avenue, Singapore 637553; School of Biological Sciences, NTU, 59 Nanyang Drive, Singapore 636921; Bioinformatics Institute, A*STAR, 30 Biopolis Street, #07-01 Matrix, Singapore 138671; Department of Biological Sciences, National University of Singapore (NUS), 16 Science Drive 4, Singapore 117558; NTU Institute of Structural Biology, Experimental Medicine Building (EMB), 59 Nanyang Drive, Level 06-01, Singapore 636921

**Keywords:** Chitin-binding domain, saturation transfer difference NMR, R&R consensus, Dosidicus gigas

## Abstract

Carbohydrate-binding proteins (CBPs) are a versatile group of proteins found in almost every organism on earth. CBPs are involved in enzymatic carbohydrate degradation and also serve as templating scaffolds in the exoskeleton of crustaceans and insects. One specific chitin-binding motif found across a wide range of arthropods’ exoskeletons is the “extended Rebers and Riddiford” consensus (R&R). However, how the R&R motif binds chitin is unclear. Here, we report the 3D structure and molecular level interactions of a chitin-binding domain (CBD-γ) located in a CBP from the beak of the jumbo squid *Dosidicus gigas*. This CBP is one of four chitin-binding proteins identified in the beak mouthpart of D. gigas and is believed to interact with chitin to form a scaffold network that is infiltrated with a second set of structural proteins during beak maturation. We used solution state NMR spectroscopy to elucidate the molecular interactions between CBD-γ and the soluble chitin derivative pentaacetyl-chitopentaose (PCP) and find that folding of this domain is triggered upon its interaction with PCP. To our knowledge, this is the first experimental 3D structure of a CBP containing the R&R consensus motif, which can be used as a template to understand in more details the role of the R&R motif found in a wide range of CBP-chitin complexes. The present structure also provides molecular information for biomimetic synthesis of graded biomaterials using aqueous-based chemistry and biopolymers.

## INTRODUCTION

Chitin is a large and insoluble structural polysaccharide consisting of a repeated sequence of N-acetyl glucosamine (NAG) monomers, which are polymerized through *β*-1,4 linkages. Chitin, after cellulose, is the most abundant biomass on earth ^**1-2**^ and is found in various organisms including the exoskeletons of insects (cuticles) and crustaceans, in the cell walls of fungi, and in various mouthparts of mollusks ^**3-5**^. The beaks of cephalopods constitute a class of chitin-containing biological materials of particular interest, owing to their high mechanical robustness ^**6**^. These mechanically-active structures are robust, wholly-organic biotools used by the animals to hunt and chew their prey. In the Humboldt squid (*Dosidicus gigas*), the beak is a biomolecular composite of proteins and nano-fibrillar chitin assembled into a mechanically graded structure, whereby the elastic modulus varies from 5 GPa at the hard rostrum to 50 MPa in the most proximal soft wing ^**7**^. The elastic modulus and hardness are intimately related to the relative content of each component, the hydration state, and the covalent cross-linking density ^**7-8**^.

In crustaceans and insects, formation of the protein-carbohydrate complex is a critical step in the biogenesis and hardening of the exoskeleton and cuticles, and has been related to the Rebers and Riddiford (R&R) Consensus ^**9**^. The R&R Consensus was originally described as G-x(8)-G-x(6)-Y-x-A-x-E-xGY-x(7)-P-x-P and was later modified to an extended R&R Consensus. The pfam sequence that represents cuticle proteins is known as pfam00379 (IPR000618) ^**9-10**^. To date, no experimental tertiary structure of a protein containing an R&R Consensus has been reported. A study combining Raman, infrared and Circular Dichroism (CD) spectroscopy measurements on a soft cuticle protein inferred that the R&R Consensus was likely to adopt a *β*-pleated sheet conformation^**11**^. Hamodrakos *et al*. recognized that the R&R consensus was homologous to bovine plasma retinol binding protein and proposed that the basic folding motif is likely to be a half-barrel of anti-parallel *β*-sheets^**12**^. They further hypothesized that the interactions with chitin are meditated via the flat surface of the protein, which is composed of aromatic side chains that would interact with the polysaccharide chains of chitin ^**13**^. To our knowledge, no experimental study has reported the structure of a CBD harbouring the R&R Consensus.

We previously demonstrated that the His-rich domains of beak proteins are involved in inter-protein cross-linking reactions *via* Michael addition between the imidazole ring of Histidine and catecholic moieties, which could either originate from post-translational modification of Tyrosine residues or from low molecular weight compounds. We postulated that the proteins may contain *β*-sheet domains ^**1**,**8**^. Integrating comprehensive transcriptomic and proteomic investigations, we subsequently identified and sequenced two sets of proteins that, together with chitin, make up the beak, namely *D. gigas* His-rich beak proteins (DgHBPs) and *D. gigas* chitin-binding beak proteins (DgCBPs) ^**1**^. From these additional findings, we correlated the stiffness of the beak with the relative protein/chitin/water content along the beak. This led us to propose that beak biogenesis occurs through a multi-step mechanism, whereby DgCBPs/chitin self-assembles into a porous scaffold that is subsequently permeated and filled by DgHBPs micro-droplets. In the final stage of the process, inter-protein cross-linking DgHBPs occurs, leading to sclerotization and hardening of the beak. Studies on recombinant DgHBPs have shown that they are mostly intrinsically disordered with a characteristic propensity for Liquid-Liquid Phase Separation (LLPS); however the structure and biophysical properties of DgCBPs remain unknown ^**14-15**^.

Here, we initially attempted to obtain the crystal structure of full length DgCBP-3, but these efforts were unsuccessful due to solubility issues. Instead, we opted to screen a wide range of shorter CBDs from DgCBPs for solubility. We were able to identify a CBD from DgCBP-3 (hereafter referred to as “CBD-γ”) that could be expressed and purified in soluble form **(Supplementary Table 1)**. We used heteronuclear Nuclear Magnetic Resonance (NMR) spectroscopy to determine the structure of CBD-γ complexed with the chitin subunit pentaacetyl-chitopentaose (PCP). Whereas CBD-γ was initially compared against a CBD of the peritrophin-44 protein, we found that the CBD-γ sequence aligns well with CBPs containing the R&R Consensus. The structure of CBD-γ was resolved in the presence and absence of unlabeled PCP. Using 2D TOCSY and NOESY NMR experiments, we showed that folding of CBD-γ is triggered by complexation with PCP chitin sub-units and derived its 3D structure, which is similar to a predicted CBD structure ^**11**^. To further understand the interaction mechanisms, PCP was docked onto the NMR-derived CBP-γ structure, using the experimental inter-molecular constraints obtained from NMR measurements. Based on NMR and molecular dynamics (MD) simulations we were able to identify specific molecular interactions between CBD-γ and chitin and propose that these contribute to the strong chitin/protein complexation during squid beak growth.

## RESULTS

### Expression, purification and biophysical characterization of CBD-

DgCBPs is one of two protein families abundant in the squid’s beak. Unlike DgHBPs, they do not vary in abundance between the soft region embedded in the buccal mass of the rostrum and the hard rostrum (**Fig. 1A**) ^**1**^ Since attempts to express full-length DgCBP-3 gave insoluble proteins, we designed constructs of the active CBDs **(Table S1)**. Soluble CBD-γ protein was obtained when fused to a glutathione S-transferase (GST) tag to increase solubility (**Fig. 1B**). The expressed recombinant fusion protein was initially purified by glutathione affinity purification followed by overnight TEV-cleavage. A second glutathione affinity purification step and subsequent gel filtration chromatography were performed to separate the cleaved GST-fusion tag from CBD-γ and eliminate the TEV protease. Thus, a 3-step purification protocol yielded a homogeneous CBD-γ protein suitable for biophysical characterization. The purity of the proteins was confirmed on a 4–20% gradient polyacrylamide gel (**Fig. S1A**). We used the sequence of the CBD-γ to search for orthologues across species ^**16**^. Surprisingly, we found that the N-terminal portion of the protein shows close similarity with the original R&R consensus previously reported for the cuticle proteins AMP1B in *H. americanus* (P81385) and the cuticle protein-14 in *L. Polyphemus* (P83354). Moreover, the C-terminus shares resemblance with the peritrophin-A domain of *A. duodenale* ^**9**^. Finally, CBD-γ was found to share high similarity, both along the C- and N-termini with two uncharacterised proteins from *O. bimaculoides* (A0A0L8H3P5) and *O. vulgaris* (XP_029637511.1) **(Fig. 1C)**.

**FIGURE 1.**
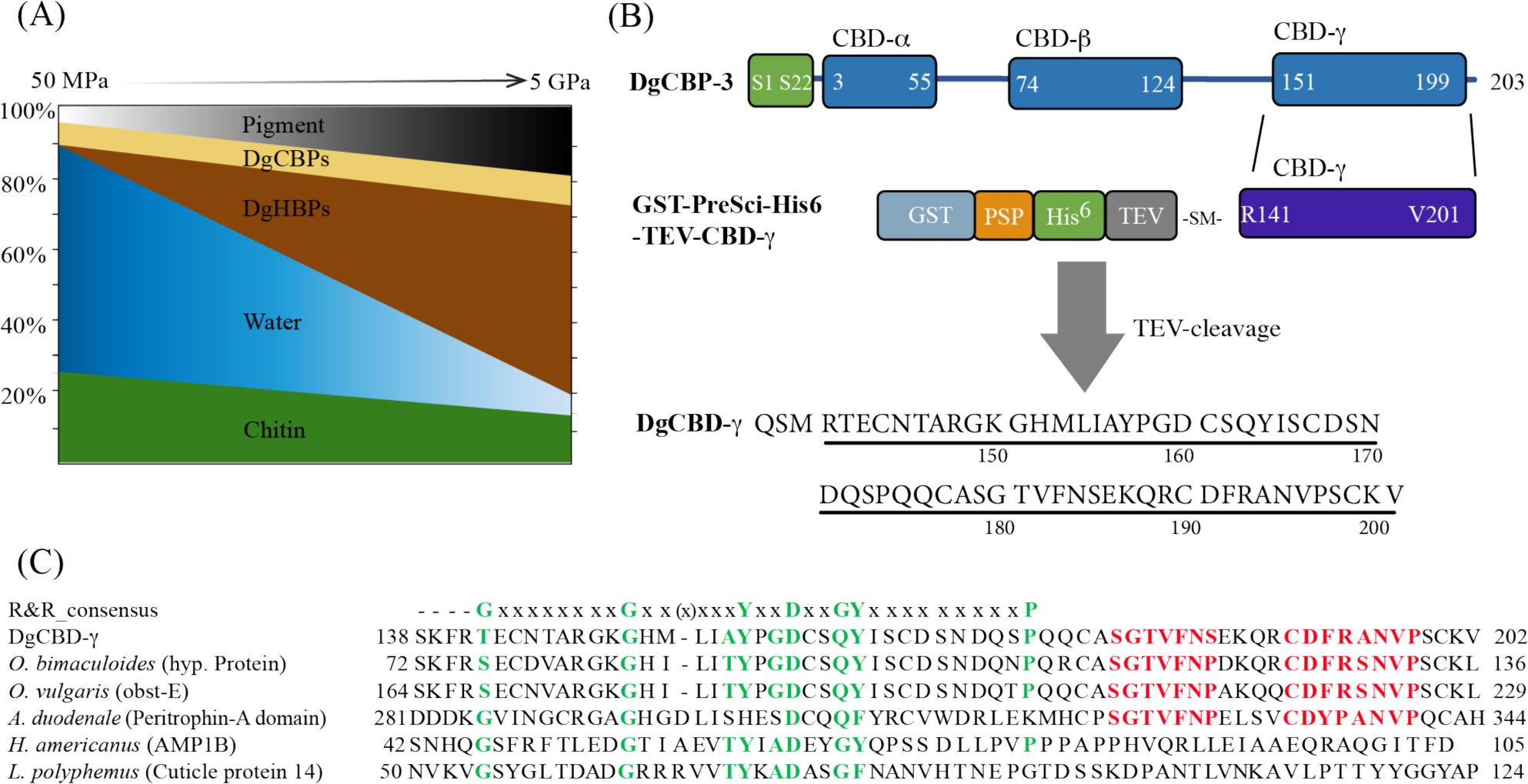
Construct design and sequence analysis of CBD-γ. **(A)** Graphical representation of the squid beak’s biomolecular gradients and their correlation with the beak’s stiffness from the soft hydrated region (elastic modulus E = 50 MPa) to the hardened, protein-rich rostrum of the beak (E = 5 GPa) (based on 1). **(B)** Construct design of CBD-γ from DgCBP. **(C)** Comparison of the sequences of CBD-γ and other proteins with the R&R consensus sequence (residues highlighted in green color), the Peritrophin-A domain (residues highlighted in red color), and with either a sequence similarity of >80% or those containing a R&R consensus sequence.

To investigate the properties of CBD-γ, biophysical characterizations were performed using far UV CD spectroscopy. β-sheets are characterized by a broad negative minima at 215 – 220 nm and a maximum at 195 nm, whereas *α*-helical structures exhibit characteristic negative minima at 208 and 222 nm and a positive maximum at 195 nm ^17^. For CBD-γ alone, the CD spectrum displayed a maximum at 192 nm and a broad minimum at 200 nm, suggesting a mostly random coil structure (**Supplementary Figure 1B**). In comparison, the CD spectrum for CBD-γ in the presence PCP displayed a shift of the broad minimum from 200 nm to 205 nm, the emergence of an additional secondary minimum at 214 nm, and a shift of the maximum from 192 nm to 197 nm, suggesting a conformational change of CBD-γ towards a structure that contains both α-helices and β-sheets.

### Saturation transfer difference (STD) NMR spectroscopy reveals that polar interactions play a dominant role in CBD-γ PCP interactions

The localization of protons in the chitin subunits of PCP onto CBD-γ was studied using 1D STD NMR spectroscopy. As the protein signals are saturated in STD NMR experiments, protons such as H1, H2, H3, NH (H2) and CH_3_ (H2) of PCP can be clearly identified from both difference and STD spectra, and used to gain insights into the CBD methods, namely epitope mapping with increasing saturation times (*t*_sat_) and dissociation constant (*K*_D_) determination with increasing PCP concentrations were employed. At the highest CBD-γ to PCP molar ratio (1:250), the STD intensities of protons H1, H2 and H3 increased with an increase in saturation times (*t*_sat_), whereas the H4 and H5 protons displayed negative STD peaks (Fig. 2A). Assessing the STD amplification factor for the five interacting protons H1, H2, H3, NH (H2) and CH_3_ (H2) of PCP at different saturation times, we calculated the STD total values as described in Eq. 4 in Materials and Methods. H1 and H2 protons exhibited higher STD total values of 21.5 and 20.8 respectively followed by NH (19.9) and CH3 (17.6) protons of H2 and H3 (11.6). (Fig. 2B). When CBD-γ was titrated with increasing concentration of PCP at *t*_*sat*_ of 2.0 s, the intensity of STD difference spectra increased, facilitating the calculation of *K*_D_ values (Fig 2B). The H1 and H2 protons exhibited the lowest *K*_D_ values followed by the CH_3_ and NH protons of H2 (Table 1). The strongest STD effect of H1 and H2 protons can be attributed to its interaction with the exposed polar residues of the protein ^19^. Hence according to group epitope mapping principle, the H1 protons are scaled to 70-100% binding and all the other protons are mapped with respect to H1 (Fig. 2D).

**Table 1.**
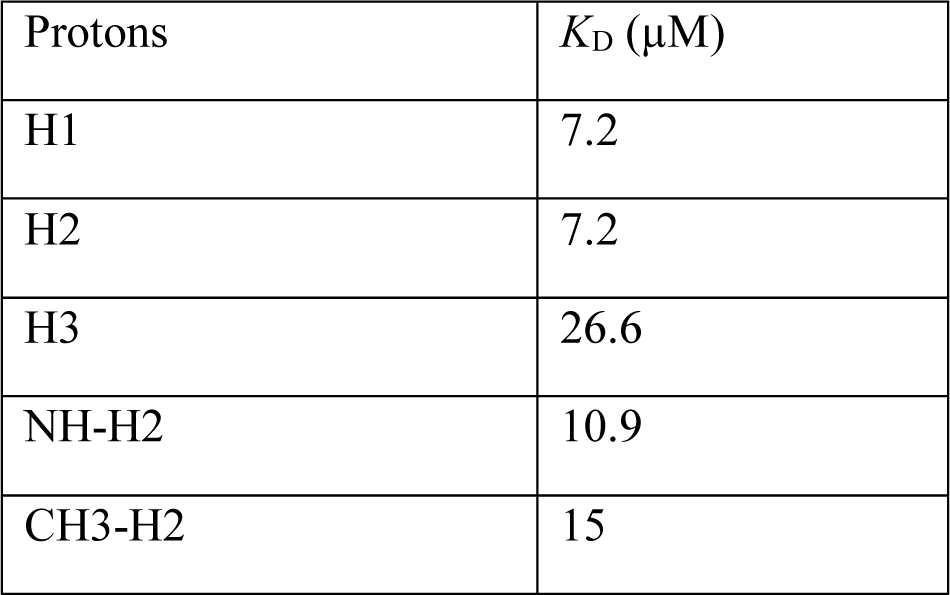
Dissociation constants (*K*_D_) of various protons of PCP in interactions with CBD-γ

| Protons | $K_D$ ($\mu\text{M}$) |
| --- | --- |
| H1 | 7.2 |
| H2 | 7.2 |
| H3 | 26.6 |
| NH-H2 | 10.9 |
| CH3-H2 | 15 |

**FIGURE 2.**
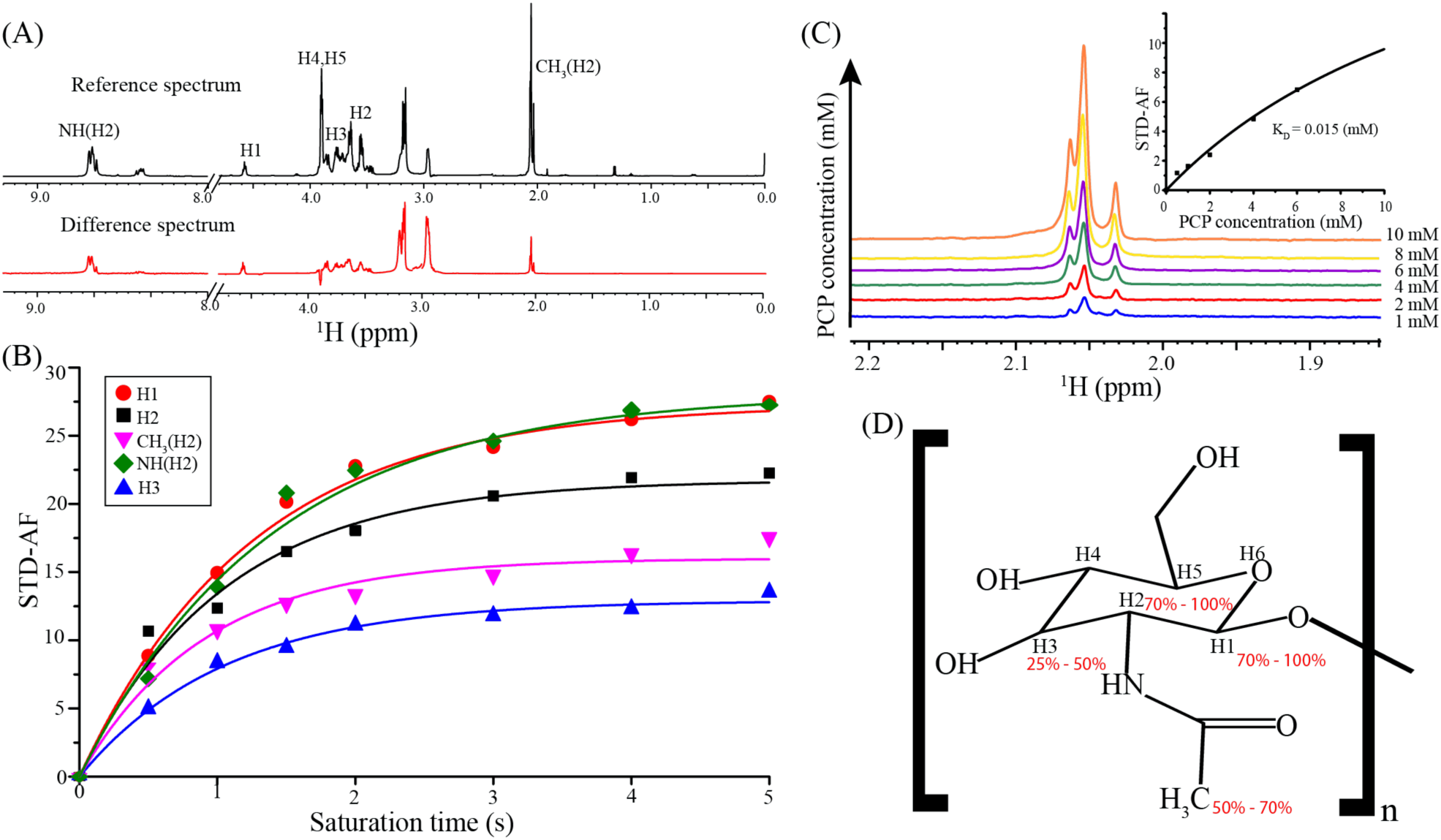
Group epitope mapping of the chitin subunit of PCP using STD-NMR. **(A)** 1D STD-reference and difference NMR spectra of CBD-γ:PCP (t_sat_ = 2.0s) with different proton resonances (H1, H2, H3, H4, H5, NH(H2) and CH_3_ (H2)) of PCP labelled. **(B)** STD amplification factor of five different proton resonances (H1, H2, H3, NH(H2) and CH_3_ (H2)) as a function of increasing saturation times (t_sat_ = 0.5 s, 1 s, 1.5 s, 2 s, 3 s, 4 s, 5 s) to calculate STD_max_ and K_st_ from the equation, STD-AF = STD_max_(1-e-^kst*tsat^). **(C)** Stacked 1D STD-difference NMR spectra of acetamide (CH_3_) region at a saturation time t_sat_ = 2.0 s, for increasing PCP concentrations. (Inset: fitting of STD amplification factor (STD-AF) against increasing PCP concentrations to obtain K_D_). **(D)** Group epitope mapping of one chitin subunit of PCP based on the STD_total_ value calculated from the equation STD_total_ = STD_max_ · K_st_. The proton that exhibits the highest STD_total_ value was defined as 100% and all other protons are normalized with respect to the highest STD_total_.

### PCP induces conformational changes in CBD-γ

The solution conformation of CBD-γ was investigated using 2D ^1^H-^1^H TOCSY and NOESY NMR experiments. Out of 61 residues (excluding prolines), 45 residues could be unambiguously assigned (**Supplementary Figure 2A**). In the next step towards obtaining the three-dimensional structure, the chemical shift deviations (CSD), which contain information about secondary structure propensity, were plotted. While negative deviations of ^1^H_α_ generally indicate helices, positive deviations indicate *β*-strands ^**20**^. The CSD plot of alpha protons of CBD-γ revealed a dynamic N-terminus with both positive and negative shifts with no clear trend apparent (**Supplementary Figure 2B**). In contrast, ^1^H_α_ at the C-terminus established a well-ordered structure with uniform positive chemical shift deviations indicating *β*-strand conformation (**Fig. S2B**). The absence of medium range NOEs and the presence of sequential HN-HN NOEs suggested that the tertiary fold of CBD-γ is dominated by extended conformations (**Supplementary Figure 2C**).

When 0.7 mM CBD-γ was added to incremental concentrations of PCP, an increase in peak intensities and minor peak broadening were observed. Notably, in 2D ^1^H-^1^H TOCSY and NOESY spectra, proton resonances from PCP increased with the saturation of the protein at a 1:1 ratio. Furthermore, H2 and H3 proton resonances of the GlcNAc subunits of PCP were found to resonate along some of the residues of CBDγ protein (**Supplementary Figure 2D**). Comparison of ^1^H-^1^H TOCSY spectra of 0.7 mM CBD-γ in the absence or presence of PCP displayed an upfield chemical shift of resonances, suggesting conformational changes of CBD-γ in the presence of PCP (**Supplementary Figure 2D**). The ^1^H-^1^H TOCSY and NOESY spectra of the equimolar CBD-γ-PCP complex were further assessed for protein structure determination. The ^1^H-^1^H TOCSY spectra of CBD-γ-PCP complex showed well resolved ^1^H_α_ peaks of CBD-γ in addition to the GlcNAc peaks from PCP (**Fig. 3A**). In addition to the 45 residues previously assigned to CBD, an additional 12 residues could be readily assigned in the presence of PCP. The chemical shift deviations of CBD-γ-PCP revealed negative deviations in the region G^159^–Q^172^ suggesting a helical propensity, followed by positive deviations between Q^176^-V^196^ indicating that the extended β-strand conformations in the C-terminal region of CBD-γ are conserved in the CBD-γ-PCP complex (**Fig. 3C**). To confirm the conformational change of CBD-γ in the presence of PCP, ^15^N-^1^H HSQC along with 3D ^15^N-HSQC TOCSY and NOESY spectra were also analyzed (**Fig. 3B**). Well-defined peaks were visible in ^1^H-^15^N HSQC spectra for CBD-γ-PCP complex over the 9.2-7.2 ppm range in ^1^H dimension (**Fig. 3B**), indicative of a well-folded protein conformation. The assignment of the protein residues was corroborated with the earlier assigned homonuclear spectra and the CSDs were also substantiated with the homonuclear spectra findings (**Fig. 3B).**

**FIGURE 3.**
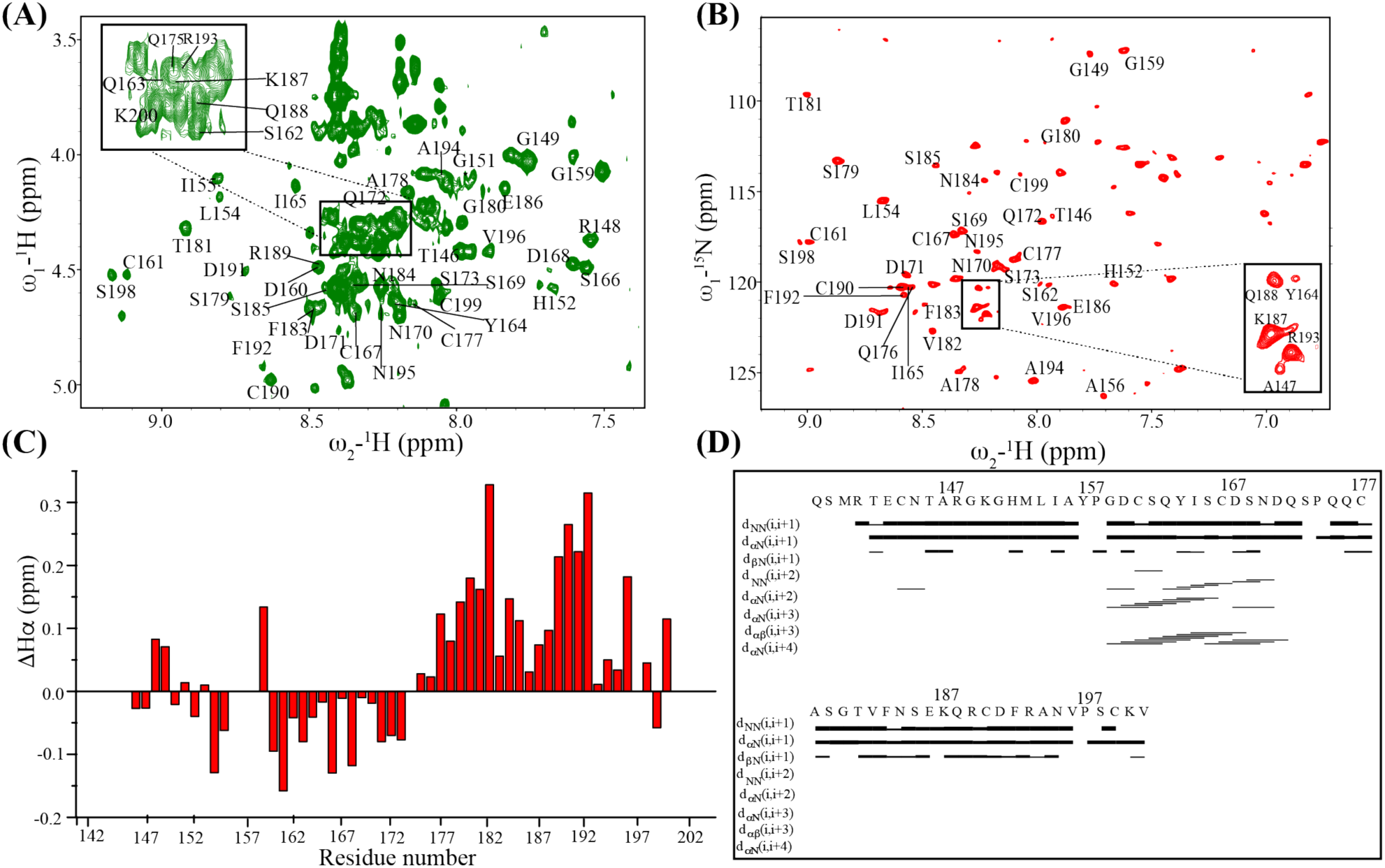
Resonance assignment of CBD-γ in presence of PCP. **(A)** Sequence specific resonance assignment in the fingerprint region of two-dimensional ^1^H-^1^H TOCSY spectrum and **(B)** 2D ^1^H-^15^N HSQC spectrum of CBD-γ in the presence of 0.7 mM PCP. **(C)** Chemical shift deviations (CSDs) from random coil of Hα of the CBD-γ - PCP complex. **(D)** Bar diagram representation of NOE connectivities used for structure calculation of the CBD-γ - PCP complex displaying the presence of medium range (*i-i* + *3* and *i-i+4*) αN from G159 – S173, indicative of helical conformations.

### Polar hydrophobic interactions stabilize the conformational changes in the CBD-γ - PCP complex

About 157 NOEs, include 27 long range NOEs, (**Table 2**) were used to determine the tertiary structure of CBD-γ. Notably, the protein was found to fold into three regions, referred to as “flexible (R^141^-P^158^), “semiflexible” (G^159^-P^174^) and “rigid” regions (Q^175^-V^201^) (**Supplementary Figure 3A**). The surface charge of CBD-γ appears to be hydrophilic and the aromatic residues are distant from each other and exposed to the solvent (**Supplementary Figure 3C and D**). The rigid, well conserved, domain is composed of three antiparallel *β*-strands with their polar side chains exposed to the solvent (**Supplementary Figure 3B**). The RMSD of the backbone residues in the rigid region was 0.436 Å.

**Table 2.**
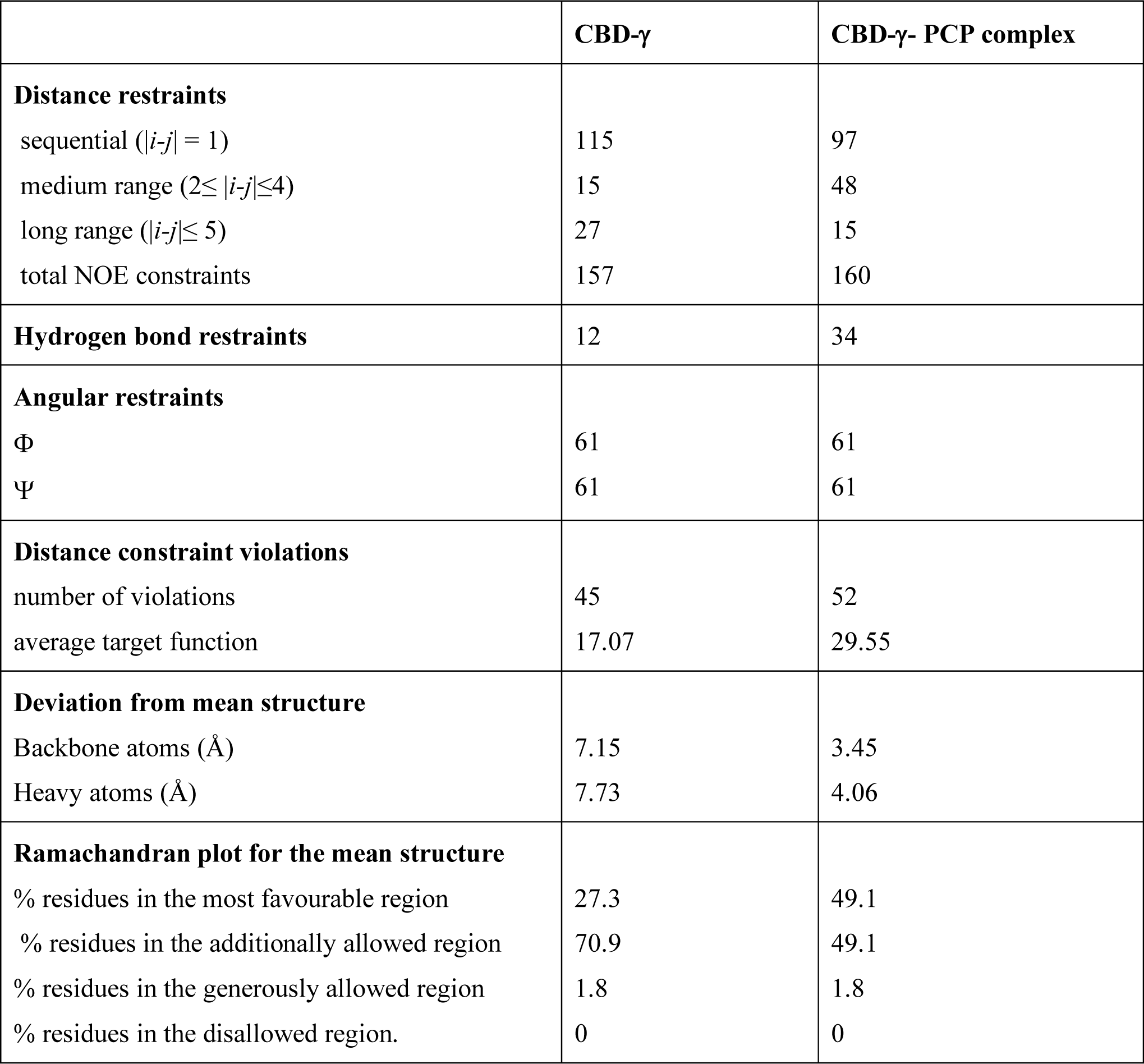
Structural statistics summary of CBD-γ and CBD-γ - PCP complex.

|  | **CBD-γ** | **CBD-γ- PCP complex** |
| --- | --- | --- |
| **Distance restraints** |  |  |
| sequential ($\vert i-j\vert = 1$) | 115 | 97 |
| medium range ($2 \leq \vert i-j\vert \leq 4$) | 15 | 48 |
| long range ($\vert i-j\vert \leq 5$) | 27 | 15 |
| total NOE constraints | 157 | 160 |
| **Hydrogen bond restraints** | 12 | 34 |
| **Angular restraints** |  |  |
| $\Phi$ | 61 | 61 |
| $\Psi$ | 61 | 61 |
| **Distance constraint violations** |  |  |
| number of violations | 45 | 52 |
| average target function | 17.07 | 29.55 |
| **Deviation from mean structure** |  |  |
| Backbone atoms ($\text{\AA}$) | 7.15 | 3.45 |
| Heavy atoms ($\text{\AA}$) | 7.73 | 4.06 |
| **Ramachandran plot for the mean structure** |  |  |
| % residues in the most favourable region | 27.3 | 49.1 |
| % residues in the additionally allowed region | 70.9 | 49.1 |
| % residues in the generously allowed region | 1.8 | 1.8 |
| % residues in the disallowed region. | 0 | 0 |

In the presence of an equimolar concentration of PCP, the semiflexible region of CBD-γ converted from a primarily unstructured conformation into an ordered α-helical conformation (**Fig. 4A**). This was further evident from the Ramachandran plot, where the number of residues in the additionally allowed region decreased with an increase in the number of residues in the most favorable region (Table 2) of the CBD-γ - PCP complex. The helix-β1 interface was strongly stabilized by hydrophobic interactions between the side chains of Y^164^ and D^168^ interfacing with residues S^179^ and Q^175^ (**Fig. 4B**). The electrostatic surface analysis of CBD-γ in the presence of PCP suggested that the conformational change, which is mediated by stabilizing hydrophobic interactions, could arise from the interactions of the aromatic rings of Y^164^ and F^183^ with the CH protons of H2 in PCP (**Fig. 4C and D).** For the determination of the tertiary structure of the complex, about 160 NOEs, including 48 medium-range and 15 long-range” NOEs were used. The RMSD of backbone residues of helical regions was calculated as 0.089 Å and that of *β-*strands as 0.474 Å. Surprisingly, despite the presence of six cysteines in the sequence, we found no evidence of disulphide bond formation in the structure.

**FIGURE 4.**
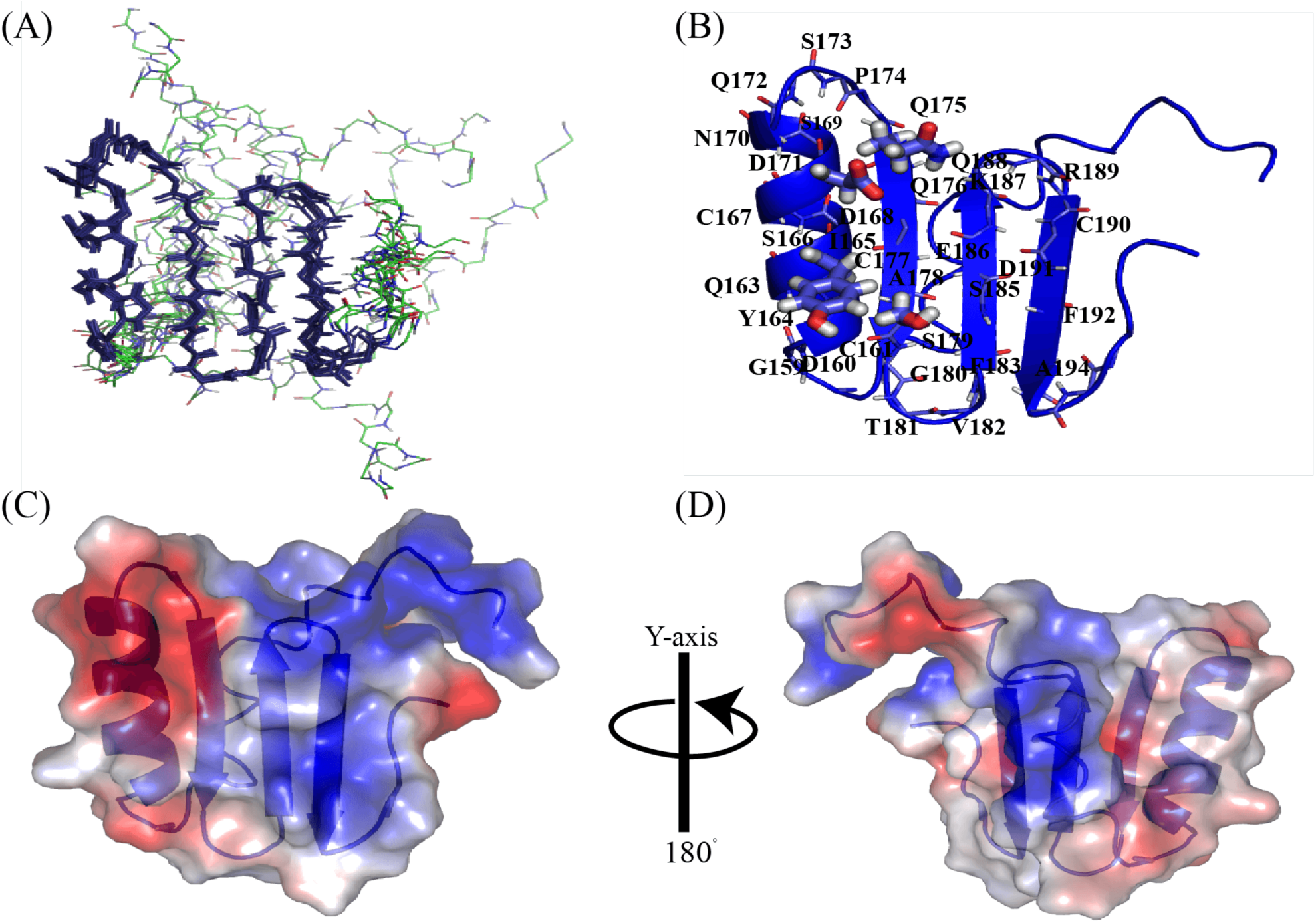
NMR structure of the CBD-γ - PCP complex. (A) An ensemble of backbone of the ten lowest energy structures of CBD-γ - PCP complex. The folded regions are represented as sticks. (B) A representative conformation of CBD-γ-PCP complex where the backbone atoms of the structurally-defined residues are marked. The side chain interactions of Y164 and D168 in the helical region interfacing with S^179^ and Q^175^ of β-sheets are represented as sticks. (C) Electrostatic surface potential map of CBD-γ-PCP complex, and (D) its 180 degrees rotated representation. The involvement of hydrophobic residues (highlighted in red color) in stabilizing interactions between helix and β1 interface can be clearly visualized.

### Mapping of PCP binding sites on CBD-γ suggest that polar interactions play a dominant role in complex formation

Chemical shift perturbation (CSP) analysis is one of the more prominent measurements to interpret binding interactions between proteins and ligands ^**21-22**^. CSP can differentiate between strong to weak binding at the residue level. Three amino acids, namely Y^164^, N^170^ and Q^176^ in the folded helical regions unambiguously exhibited strong perturbation in the presence of an equimolar concentration of PCP. Additionally, polar residues from *β*-sheets displayed weaker perturbations (**Fig. 5A**).

**FIGURE 5.**
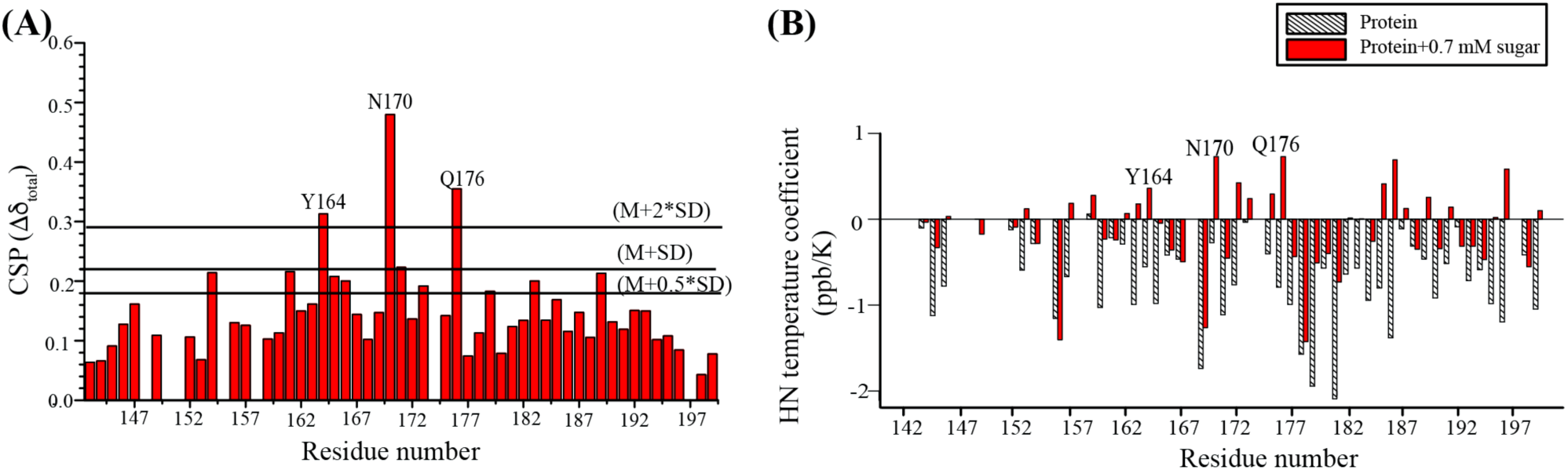
CSP and Amide temperature coeffiecients of CBDγ. **(A)** CSP plot of CBD-γ - PCP complex plotted against residue number. The residues of CBD-γ that show stronger perturbations (above the mean + 2 x standard deviation (SD) line) are marked. **(B)** Amide temperature coefficients of CBD-γ and CBD-γ - PCP complex plotted against residue number. The residues with stronger CSP show an increase in hydrogen bonding efficiency as marked.

However, the weaker perturbations for residues such as S^179^, T^181^ and F^183^ continued to evolve with increase in PCP concentrations. At 1.2 mM PCP concentration, the chemical shifts of these residues suggest exchange between the free and bound forms (**Supplementary Figure 4A**). Chemical shift perturbations of Y^164^ and F^183^ were further supported by calculating the amide temperature coefficients of CBD-γ in the absence and presence of PCP. While the NH temperature coefficient in the absence of PCP showed continuous negative values in the range of -0.1 ppm / K and −2.1 ppm / K, less negative and more positive coefficients were observed in the presence of PCP **(Fig 5B black/white bars)**. In contrast, the amide temperature coefficients of CBD-γ in the presence of PCP returned positive coefficients, including the same amino acids (Y^164^, N^170^ and Q^176^) that previously exhibited the strongest perturbations (**Fig. 5B, red bars**). When the perturbed residues were mapped onto the tertiary structure of CBD-γ two major interactions that mediate binding could be identified (**Supplementary Figure 4B**). First are the hydrophilic polar interactions between the electron-rich hydroxyl groups of the protein’s polar residues (S^166^, N^170^, S^173^, S^179^) and the electron-deficient acetamide groups of the carbohydrate (**Supplementary Figure 4B**). Second, the electronegative aromatic (π) rings of Y^164^ and F^183^ interacted with the electropositive CH groups of the H2 protons bringing about a stronger hydrophobic effect to the binding. To confirm the above findings, we generated alanine mutant of CBD-γ (Y^164^→A, N^170^→A, Q^176^→A) and conducted a binding assay with chitin magnetic beads followed by analytical FPLC. The experiment showed that the mutant did not have any activity to bind to chitin (**Supplementary Figure 5**).

The CSPs recorded experimentally by NMR were used to model the interactions between CBD-γ and PCP through computational docking. The strong chemical shifts associated with specific residues (Y^164^, N^170^, Q^176^) in CBD-γ were used as restraints to computationally dock PCP. The 10 docked poses were subjected to MD simulations and in 7 poses, the PCP dissociated from the protein. In the remaining 3 poses stable bound conformations were seen during the MD simulations and all 3 systems showed a similar binding mode and inter-molecular contacts. In these 3 cases, in accord with the NMR observations, PCP was stably bound in a groove formed by the β-sheet and the helix, lying orthogonal to the direction of the helix (involving N_163_, Y^164^, S^166^, D^168^, N^170^, N^172^, F^183^, S^185^, R^193^ residues, **Fig 6A**). Energetics analysis of the complex during the MD simulations suggests that the complexation is driven primarily by Van der Waals interactions **(Fig 6A and B)**. While the electrostatic component is large, the desolvation penalty dominates. To further collate insights about the interactions, alanine mutants CBD-γ (Y^164^→A, N^170^→A, Q^176^→A) were generated computationally and analyzed. The final conformation sampled in one of the 3 MD simulations (which showed stable complexes) was taken. The interacting residues were mutated to Alanine and the resulting complex subject to 100 ns of MD simulations. Within 40 ns, the PCP dissociated from the mutant CBD-γ **(Fig 6C, inset plot)** and in parallel, the protein exhibited substantial unfolding **(Fig. 6C**, inset structure figure). The distance distribution plot **(Fig. 6C)** showed a single peak at ∼4 Å for the wild-type. However, for the mutant 3 peaks were observed. The first peak at *ca.* 6 Å corresponded to the initial conformation (up to 40 ns, **Fig. 6C** inset plot); the second peak observed at *ca.* 8 Å corresponded to the initial unbinding (50-70 ns) followed by a complete dissociation observed towards the end of the MD trajectory (with third peak at *ca.* 11.8 Å). Loss in PCP interaction was also accompanied by order to disorder transitions in CBD-γ (**Fig. 6C** inset, structure shown in cartoon representation). These findings underscore the importance of these residues in establishing key interactions with PCP, as also corroborated experimentally with the chitin bead binding assay (**Supplementary Figure 5**).

**FIGURE 6.**
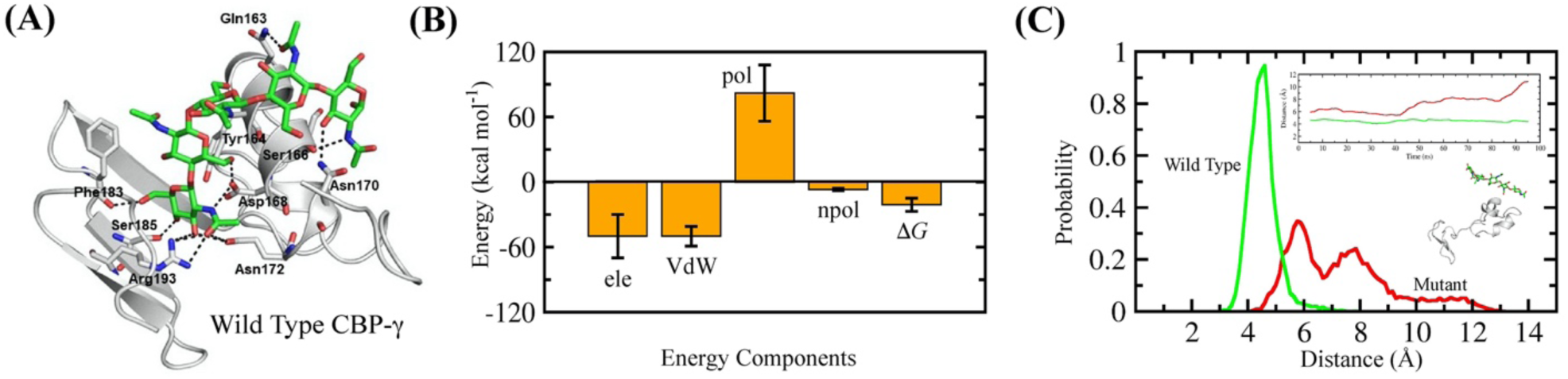
Molecular Dynamics of CBD-γ. **(A)** CBD-γ grey cartoon in complex with PCP (green sticks). The key interacting residues are highlighted and shown as sticks. **(B)** Energy components of the total free energy of interaction between CBD-γ and PCP during the MD simulations, Ele: electrostatics, VdW: Van der Waals, pol: polar, npol: non-polar contribution to solvation free energy, **Δ***G*: net binding free energy. **(C)** Distribution plot depicting the distance between center of mass (COM) of interacting residues of CBD-γ (wild type: green and mutant: red) with PCP. **Insets**: the time dependent changes in the averaged distance between COM of interacting residues of CBD-γ and PCP over the MD trajectory and a cartoon representation of the complex illustrating the dissociation of PCP and the destabilized mutant CBD-γ structure.

## DISCUSSION

DgCBPs are an important group of proteins contributing towards the structural integrity of squid beaks. The current study focused on the tertiary structure determination using solution NMR of the chitin-binding domain CBD-γ in complex with chitin subunits (PCP). Our data show that structure of CBD-γ exhibits a relatively flexible N-terminal domain followed by a partially flexible central domain and a rigid C-terminal domain with a three-stranded β-sheet. The addition of PCP initiated a ligand-induced conformational change leading to the formation of an α-helix from the partially flexible central domain of CBD-γ. These observations are also recapitulated when the PCP is removed from the complex between CBD-γ and PCP and is subjected to MD simulations. The resulting apo structure of CBD-γ undergoes structural destabilization. We found that the partially flexible region of CBD-γ contains an extended R&R Consensus RR-2 which plays a critical role in binding to chitin during the formation of insect cuticles ^**23**^. Our study experimentally establishes that the conformational change of the protein is triggered by polar residues from the chito-oligomer, which actively interact with the polar residues protruding from the α-helix and the β1-strand of CBD-γ. Our hypothesis is supported by the change of the protein surface net charge and the involvement of three polar residues, Y_164_, N_170_, and Q_176_. While the aromatic ring protons could interact well with the saccharide ring CH groups of PCP, the polar residues are likely to bind to the acetamide group of chitin side chains. In agreement with the experimental studies, MD analysis indicates that along with the aromatic residues, polar residues add specificity to the interaction. The latter act as anchoring residues at the edges of the putative binding surface. The retained interactions identified from the MD analysis are also in agreement with the NMR observations. Computationally mutating the key interacting residues of CBD-γ and subjecting the mutant complex to MD simulations results in dissociation of the PCP and destabilization of the CBD-γ. The conformational switch also explains the almost irreversible binding that has been reported between chitin and proteins found in the squid beak. Thus, we suggest that these findings might be directly relevant to other chitin/protein biological structures of interest such as the exoskeleton of crustaceans or insects ^**1**^.

Using STD-NMR, we identified the carbohydrate side chains involved in complex formation and confirmed that the carbohydrate protons H1, H2, H3, NH-H2 and CH3-H2 –all positioned on one side of the carbohydrate– are interacting with CBD-γ. The remaining protons on the carbohydrates, on the other hand, are not exposed to the protein. All protons show *K*_D_ values in the low micromolar range, indicating a strong affinity towards the protein. Whereas previous studies on insect cuticle proteins containing the R&R consensus predicted structural features of CBMs based on homology modelling ^**13**^, experimental determination of R&R-containing domains has previously not been reported. Combining NMR and MD simulations, we note that the observed binding mechanism mediated via polar residues of CBD-γ to PCP is similar to the binding of other chitin-binding proteins, such as tachycitin or hevein, although they do not share the same structure ^**13, 24 25**^.

In conclusion, our results provide the first experimentally determined structure of a domain harboring the R&R consensus sequence, which plays a central role in stabilizing chitin/protein load-bearing structures identified in a wide range of phyla ^**26**^. Our approach and findings may inform research conducted on other structural chitin-binding domains across species, including other cephalopods, crustaceans, and insects ^**27-29**^. Further, the presented binding mechanism is also relevant to studies on other non-structural chitin-binding domains, such as enzymatic chitinases ^**30**^.

## EXPERIMENTAL PROCEDURES

### Cloning and expression of CBD-γ

Residue numbering of DgCBP-3^R141-V201^ (hereafter referred to as CBD-γ) is based on UniProt ID A0A0G2UHG5. The gene encoding CBD-γ was submitted to the Protein Production Platform at the School of Biological Sciences, Nanyang Technological University, Singapore. Cloning was performed using ligation-independent cloning (LIC) technology in *E. coli* and the vector used was pNIC-GST ^**31**^. The construct of CBD-γ fused to a glutathione S-transferase (GST) tag to increase solubility, was amplified using PCR with the primers R141-TACTTCCAATCCATGCGCACCGAGTGCAATACTG-V201 and V201-TATC CACCTTTACTGTCAGACCTTACAGCTTGGC ACAT-R141 and expressed in BL21(DE3) Rosetta T1R. The target protein sequence was fused to the GST protein N-terminal followed by a hexa-His-tag and a TEV-protease cleavage site. The transformed cells were streaked onto Luria-Bertani (LB) agar plates with Chloramphenicol (26 µg mL^-1^) and Kanamycin resistance (50 µg mL^-1^). Single colony transformants were expressed with 2 × LB media and protein expression was induced with 0.3 mM IPTG for 18 hours at 17°C.

### Purification of CBD-γ

A gene of CBD-γ was overexpressed in *Escherichia coli* BL21(DE3) in either LB media or in M9 minimal media containing ^15^N-labeled NH_4_Cl. Cells were harvested using centrifugation (4,000 × g, JLA-8.1000, Beckman) for 20 minutes (Avanti JXN-30 Series, Beckman). Cell pellets were collected and suspended in a washing buffer (20 mM Na HEPES, pH 6.5, 500 mM NaCl), supplemented with cOmplete™ Protease Inhibitor Cocktail (1697498001, Roche). Cells were lysed using high sheer fluid processing (LM20 Microfluidizer, Microfluidics) at 25,000 psi. The lysate was clarified by centrifugation (20,000 × g, JLA-25.50, Beckman) and the soluble fraction separated from the cell debris. A volume of 3 mL of Glutathione Sepharose 4B resin (17075605, GE Life Sciences) was added to the soluble fraction and incubated for a minimum of 6 hours at 4°C. Unbound proteins were removed by washing the beads with 10 × the column volume (CV) and 5 mL of washing buffer was added to the beads. Elution of the target protein was initiated by adding TEV protease to the solution and incubated overnight at 4°C. The supernatant was separated from the resin and injected in Superdex 75 16/600 column pre-equilibrated with the washing buffer and eluted fractions were monitored using an NGC Quest™ 10 Plus Chromatography System (Bio-Rad). The fractions were collected and concentrated to a final concentration of 700 µM using Vivaspin sample concentrators (GE Healthcare; cut-off: 3 kDa). The protein purity was analysed by SDS-PAGE using 4–15% Mini-PROTEAN® TGX™ Precast Protein Gels (Bio-Rad).

### Circular Dichroism (CD) experiments

The far-UV CD spectra of CBD-γ and CBD - PCP were measured using a Chirascan spectropolarimeter (Model 420, AVIV Biomedical Inc.), using Quartz supracil cells (0.2 mm path length; Hellma Analytics). The spectra were recorded in triplicates at 18°C, at wavelengths ranging from 240 nm to 190 nm using 1 nm increments and a 1 nm bandwidth, with instrument dynode voltage less than 600 nm. The baseline was acquired using the protein sample buffer. All spectra were smoothed using the Savitzky-Golay filter using MATLAB and the acquired spectra adjusted by subtracting the smoothed, mean averaged baseline spectrum from the smoothed, mean averaged protein sample spectrum ^**32**^.

### STD-NMR experiments

Saturation Transfer Difference (STD) NMR titration assays were carried out using a concentration of 50 µM of CBD-□ protein at 4°C. PCP at ratios ranging from 12.5 µM to 250 □M were added to the protein. The standard *stddiffesgp.3* pulse sequence was used with a saturation length of 50 ms at a power of 50 db. The *on*- and *off*-resonance irradiations were applied at a chemical shift of −1.2 ppm and 40 ppm, respectively with saturation time (t_sat_) of 2.0 s. All spectra were processed using the *stdsplit* macro in Topspin (Bruker). As defined by *Mayer & Meyer*, the STD amplification factor was calculated with following equation **(Eq. 1)** where *I*_0_-*I*_sat_ is the intensity of difference spectra and I_0_ is the intensity of the reference spectra ^**33**^:

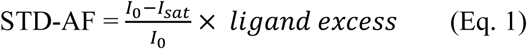

The *K*_D_ of given protons was then obtained by fitting STD amplification factor against ligand (PCP) concentration using a non-linear fitting to ^**34**^:

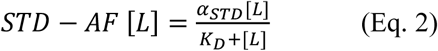

Group epitope mapping was conducted at the highest concentration of PCP at a protein-to-carbohydrate molar ratio of 1:250 at various saturation times (D20) ranging from 0.5 s to 5.0 s. The D1 relaxation delay was set to 5.5 s. The STD-AF of different protons such as H1, H2, H3, NH (H2) and CH_3_ (H2) were derived using the above-mentioned equation **(Eq. 1)** and STD_max_ and K_st_ were calculated by fitting the STD-AF against saturation times using the following equation:

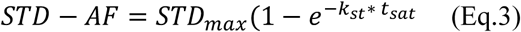

Subsequently, STD_total_ was calculated using the following equation:

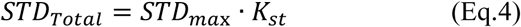

Lastly, the proton that recorded the highest value for STD_total_ was considered as binding 100% to PCP in accordance with the epitope mapping rule in order to derive the binding contribution of all the remaining protons of CBD-γ towards PCP ^**35**^.

### NMR Experiments and Structure Calculations

All NMR experiments were carried out in Bruker 700 MHz or 800 MHz spectrometers equipped with cryoprobes at 4°C. About 0.7 mM of CBD-γ stored in analysis buffer (20 mM Na HEPES, pH 6.5, 500 mM NaCl) were supplemented with 10% (v/v) D2O for deuterium lock and DSS (2,2-dimethyl-2-silapentanesulfonic acid) for signal reference. Two-dimensional ^1^H-^1^H TOCSY and ^1^H-^1^H NOESY spectra of CBD-γ were acquired with 80 ms and 200 ms mixing times respectively. There were 512 increments in F1 and 2048 in F2 axes with the number of scans set to 32 for TOCSY and 64 for NOESY experiments. To measure the effect of the presence of chitin on the stability of CBD-γ, increasing concentrations of PCP (0.35 mM, 0.7 mM, 1.2 mM) was added, and a series of ^1^H-^1^H TOCSY and ^1^H-^1^H NOESY were recorded after each addition.

All NMR data were processed using TOPSPIN (Bruker) and analyzed using NMR-FAM Sparky programs. NMR structures were calculated using the CYANA 2.1 program. The intensities of NOE cross-peaks were classified as strong, medium, and weak and translated to upper bound distance limits of 2.5, 3.5, and 5.0 Å. The dihedral Φ and Ψ angles were constrained between −120° and −30° and between −120° and 120° as suggested in the CYANA program files. Of the 100 structures generated, the 10 lowest energy structures were used for analysis. Structure representations were made using PyMOL Molecular Graphics System ^**36**^. Electrostatics calculations were carried out using APBS plugin available in PyMOL. Procheck was used to validate the calculated NMR structures ^**36-37**^.

### Chemical shift perturbation and amide temperature coefficient NMR studies

The chemical shift perturbations were calculated using ^15^N labeled CBD-γ A series of two-dimensional ^1^H-^15^N HSQC spectra were acquired with 0.7 mM CBD-γ and incrementing concentrations of PCP. The perturbations of amino acid residues of CBD-γ upon binding to PCP were determined using the formula:

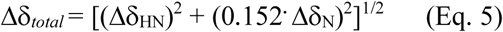

The weighing factor 0.152 was calculated as the ratio of the range of chemical shift dispersion of HN to N ^38^. The CSPs were categorized as strong if Δδ_*total*_ > mean +2σ, medium if Δδ_*total*_ > mean + 1σ and weak if Δδ_*total*_ > mean + 0.5σ. Three-dimensional ^15^N NOESY HSQC and ^15^N TOCSY HSQC were also acquired at CBD-γ - PCP (1:1) concentration to facilitate tertiary structure refinement. Amide temperature coefficients were also determined by recording 2D ^1^H-^15^N HSQC spectra of either CBD-γ or CBD-γ - PCP (1:1) at 5°C, 10°C, 15°C, 20°C and 15°C. Amide proton chemical shift deviations were fitted linearly against temperature and the temperature coefficients were calculated as σδHN/ΔT (ppb K^-1^) ^39^.

### Computational Docking and Molecular Dynamics Simulations

The first model of the NMR derived ensemble of 10 models of CBD-γwas chosen for docking and subsequent studies. It was initially subjected to energy minimization for the docking studies using the ff14SB forcefield ^**40**^ in the program AMBER18 ^**41**^. The 3D structure of PCP was taken from ChemSpider (CSID27445378). To construct models of the complex between CBD-γ and PCP, it was necessary to generate multiple conformations of PCP as it is highly flexible. For this, we subjected it to energy minimization and molecular dynamics (MD) simulations using the ff14SB forcefield in AMBER18; its parameters were generated using the Antechamber module of the program AMBER. The simulated conformations of PCP were clustered using the *kclust* program available as part of MMTSB-tools^**42**^. The top 3 conformations sampled from the highest populated cluster were then used for computational docking to CBD-γ. The PCP conformations extracted were prepared using *LigPrep* (Schrodinger 9.0) ^43^ and were docked to the minimized model of CBD-γ (see above) using the *Glide* module of the Schrodinger program following established standard protocols ^44^. 10 conformations of the docked complex were identified in the docking and these were all subject to 100 ns MD simulations each, using the standard protocol in AMBER18 ^41^. The sampled conformations were subject to clustering based on RMSD using the *kclust* module of MMTSB-tools ^42^. The structure with RMSD closest to the centroid of the most populated cluster was chosen for analysis. Analysis of simulations were carried out using *ptraj* module in AMBER. The binding energetics for the complex was calculated using MMPBSA methodology. A total of 8000 structures extracted from the last 80 ns of the equilibrated trajectory were used for the analysis. The simulated MD trajectories were viewed in VMD ^45^ and the figures were generated using PyMOL ^36^.

## Acknowledgements

This research was funded by the Singapore Ministry of Education (MOE) through an Academic Research Fund (AcRF) Tier 2 grant (Grant MOE2015-T2-1-062). We also acknowledge financial support from the US Office of Naval Research – Global (ONR-G), grant no. N62909-17-1-2045. The authors thank the A*STAR Computing Resource Centre for access to computing facilities.

## FOOTNOTES

Supporting Information include: General remarks, additional figures S1-S5, and table S1. Coordinates for the CBD-γ-PCP complex have been uploaded to the Protein Data Bank and Biological Magnetic Resonance Data Bank: PDB # 6L9Y, BMRD # 36298.

## The abbreviations used are

PCP: Pentaacetyl-chitopentaose;
CBD: Chitin-binding domain;
STD-NMR: Saturation transfer difference-NMR;
NOESY: nuclear overhauser enhancement spectroscopy;
TOCSY: Total Correlated Spectroscopy,
CMB: Chitin Magnetic Beads

## TABLE OF CONTENTS ARTWORK

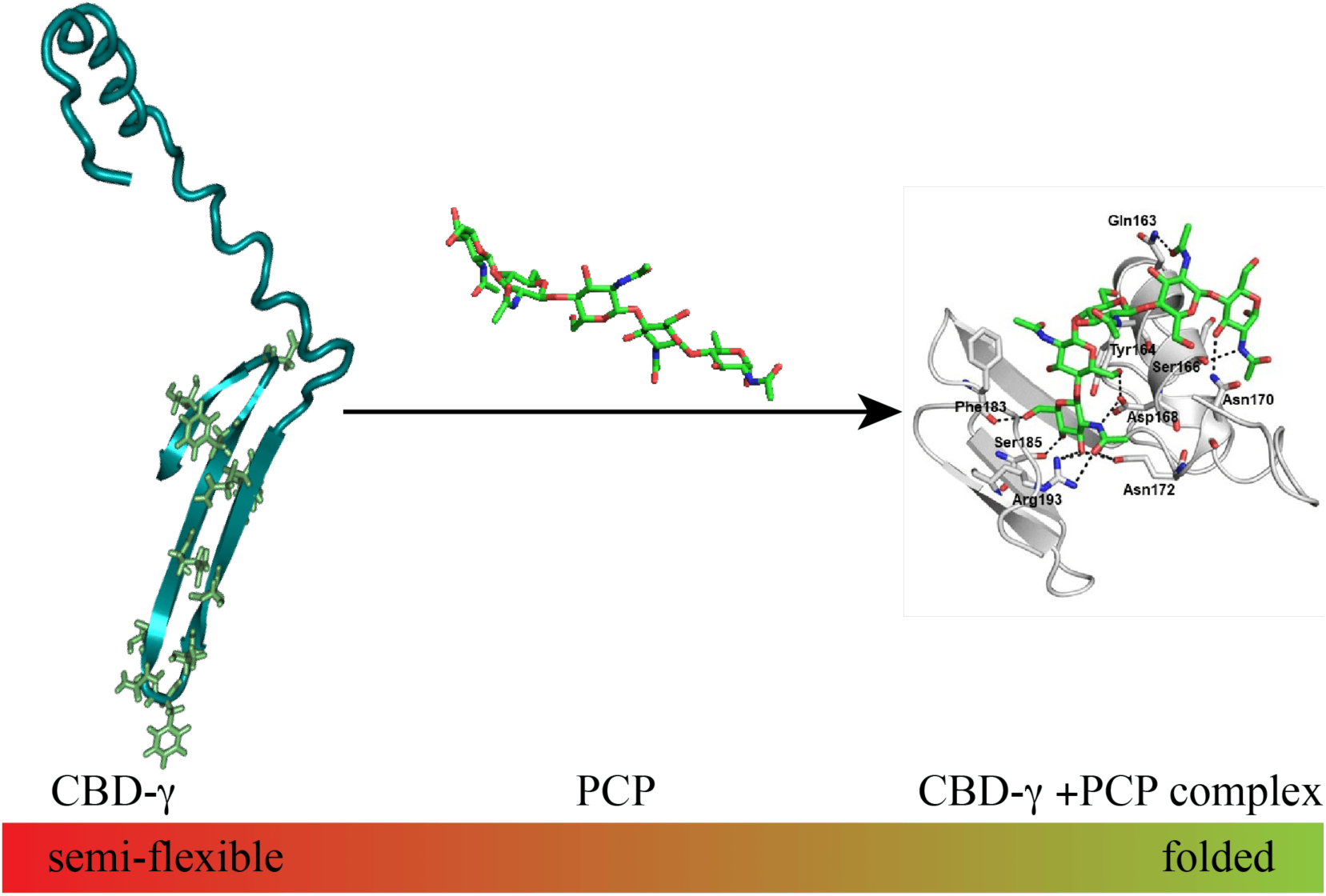

## REFERENCES

1. Tan, Y.; Hoon, S.; Guerette, P. A.; Wei, W.; Ghadban, A.; Hao, C.; Miserez, A.; Waite, J. H., Infiltration of chitin by protein coacervates defines the squid beak mechanical gradient. Nat Chem Biol 2015, 11 (7), 488–95.

2. Tharanathan, R. N.; Kittur, F. S., Chitin--the undisputed biomolecule of great potential. Crit Rev Food Sci Nutr 2003, 43 (1), 61–87.

3. Hackman, R. H., The integument of arthropoda. In Chemical Zoology, Florkin, N. M.; Scheer, B. T., Eds. Academic Press: New York, 1971; Vol. 6, pp 1–62.

4. Latge, J. P., The cell wall: a carbohydrate armour for the fungal cell. Mol Microbiol 2007, 66 (2), 279–90.

5. Peters, W., Occurence of chitin in mollusca. Comp Biochem Physiol B 1972, 41, 541–550.

6. Miserez, A.; Li, Y.; Waite, J. H.; Zok, F., Jumbo squid beaks: inspiration for design of robust organic composites. Acta Biomater 2007, 3 (1), 139–49.

7. Miserez, A.; Schneberk, T.; Sun, C.; Zok, F. W.; Waite, J. H., The transition from stiff to compliant materials in squid beaks. Science 2008, 319 (5871), 1816–9.

8. Miserez, A.; Rubin, D.; Waite, J. H., Cross-linking chemistry of squid beak. J Biol Chem 2010, 285 (49), 38115–24.

9. Rebers, J. E.; Riddiford, L. M., Structure and expression of a Manduca sexta larval cuticle gene homologous to Drosophila cuticle genes. J Mol Biol 1988, 203 (2), 411–23.

10. Rebers, J. E.; Willis, J. H., A conserved domain in arthropod cuticular proteins binds chitin. Insect Biochem Mol Biol 2001, 31 (11), 1083–93.

11. Iconomidou, V. A.; Chryssikos, G. D.; Gionis, V.; Willis, J. H.; Hamodrakas, S. J., “Soft”-cuticle protein secondary structure as revealed by FT-Raman, ATR FT-IR and CD spectroscopy. Insect Biochem Mol Biol 2001, 31 (9), 877–85.

12. Hamodrakas, S. J.; Willis, J. H.; Iconomidou, V. A., A Structural Model of the Chitin-Binding Domain of Cuticle Proteins. Insect Biochemistry and Molecular Biology 2002, 32, 1577–1583.

13. Hamodrakas, S. J.; Willis, J. H.; Iconomidou, V. A., A structural model of the chitin-binding domain of cuticle proteins. Insect Biochem Mol Biol 2002, 32 (11), 1577–83.

14. Cai, H.; Gabryelczyk, B.; Manimekalai, M. S. S.; Gruber, G.; Salentinig, S.; Miserez, A., Self-coacervation of modular squid beak proteins - a comparative study. Soft Matter 2017, 13 (42), 7740–7752.

15. Le Ferrand, H.; Duchamp, M.; Gabryelczyk, B.; Cai, H.; Miserez, A., Time-Resolved Observations of Liquid–Liquid Phase Separation at the Nanoscale Using in Situ Liquid Transmission Electron Microscopy. J. Am. Chem. Soc. 2019, 141 (17), 7202–7210.

16. Madden, T., The BLAST Sequence Analysis Tool. In The NCBI Handbook, McEntyre, J.; Ostell, J., Eds. National Center for Biotechnology Information (US): Bethesda (MD), 2002.

17. Greenfield, N. J., Using circular dichroism spectra to estimate protein secondary structure. Nat Protoc 2009, 1 (6), 2876–2890.

18. Kasaai, M. R., Determination of the degree of N-acetylation for chitin and chitosan by various NMR spectroscopy techniques: A review. Carbohydrate Polymers 2010, 79 (4), 801–810.

19. Wimmerová, M.; Kozmon, S.; Nečasová, I.; Mishra, S. K.; Komárek, J.; Koča, J., Stacking Interactions between Carbohydrate and Protein Quantified by Combination of Theoretical and Experimental Methods. PLOS ONE 2012, 7 (10), e46032.

20. Wishart, D. S.; Sykes, B. D.; Richards, F. M., The chemical shift index: a fast and simple method for the assignment of protein secondary structure through NMR spectroscopy. Biochemistry 1992, 31 (6), 1647–51.

21. Breton, R. C.; Reynolds, W. F., Using NMR to identify and characterize natural products. Nat Prod Rep 2013, 30 (4), 501–24.

22. Williamson, M. P., Chemical Shift Perturbation. In Modern Magnetic Resonance, Graham, A. W., Ed. Springer, Cham: London, 2017; Vol. 1, pp 1–19.

23. Andersen, S. O., Amino acid sequence studies on endocuticular proteins from the desert locust, Schistocerca gregaria. Insect Biochem Mol Biol 1998, 28, 421–434.

24. Nicholls, A.; Sharp, K. A.; Honig, B., Protein folding and association: insights from the interfacial and thermodynamic properties of hydrocarbons. Proteins 1991, 11 (4), 281–96.

25. Asensio, J. L.; Arda, A.; Canada, F. J.; Jimenez-Barbero, J., Carbohydrate-aromatic interactions. Acc Chem Res 2013, 46 (4), 946–54.

26. Liu, J.; Li, S.; Li, W.; Peng, L.; Chen, Z.; Xiao, Y.; Guo, H.; Zhang, J.; Cheng, T.; Goldsmith, M. R.; Arunkumar, K. P.; Xia, Q.; Mita, K., Genome-wide annotation and comparative analysis of cuticular protein genes in the noctuid pest Spodoptera litura. Insect Biochem Mol Biol 2019, 110, 90–97.

27. Stahl, A. L.; Charlton-Perkins, M.; Buschbeck, E. K.; Cook, T. A., The cuticular nature of corneal lenses in Drosophila melanogaster. Dev Genes Evol 2017, 227 (4), 271–278.

28. Zhou, Y.; Badgett, M. J.; Orlando, R.; Willis, J. H., Proteomics reveals localization of cuticular proteins in Anopheles gambiae. Insect Biochem Mol Biol 2019, 104, 91–105.

29. Abehsera, S.; Weil, S.; Manor, R.; Sagi, A., The search for proteins involved in the formation of crustacean cuticular structures. Hydrobiologia 2018, 825 (1), 29–45.

30. Tanaka, H.; Akutsu, H.; Yabuta, I.; Hara, M.; Sugimoto, H.; Ikegami, T.; Watanabe, T.; Fujiwara, T., A novel chitin-binding mode of the chitin-binding domain of chitinase A1 from Bacillus circulans WL-12 revealed by solid-state NMR. FEBS Lett 2018, 592 (18), 3173–3182.

31. Structural Genomics, C.; China Structural Genomics, C.; Northeast Structural Genomics, C.; Graslund, S.; Nordlund, P.; Weigelt, J.; Hallberg, B. M.; Bray, J.; Gileadi, O.; Knapp, S.; Oppermann, U.; Arrowsmith, C.; Hui, R.; Ming, J.; dhe-Paganon, S.; Park, H. W.; Savchenko, A.; Yee, A.; Edwards, A.; Vincentelli, R.; Cambillau, C.; Kim, R.; Kim, S. H.; Rao, Z.; Shi, Y.; Terwilliger, T. C.; Kim, C. Y.; Hung, L. W.; Waldo, G. S.; Peleg, Y.; Albeck, S.; Unger, T.; Dym, O.; Prilusky, J.; Sussman, J. L.; Stevens, R. C.; Lesley, S. A.; Wilson, I. A.; Joachimiak, A.; Collart, F.; Dementieva, I.; Donnelly, M. I.; Eschenfeldt, W. H.; Kim, Y.; Stols, L.; Wu, R.; Zhou, M.; Burley, S. K.; Emtage, J. S.; Sauder, J. M.; Thompson, D.; Bain, K.; Luz, J.; Gheyi, T.; Zhang, F.; Atwell, S.; Almo, S. C.; Bonanno, J. B.; Fiser, A.; Swaminathan, S.; Studier, F. W.; Chance, M. R.; Sali, A.; Acton, T. B.; Xiao, R.; Zhao, L.; Ma, L. C.; Hunt, J. F.; Tong, L.; Cunningham, K.; Inouye, M.; Anderson, S.; Janjua, H.; Shastry, R.; Ho, C. K.; Wang, D.; Wang, H.; Jiang, M.; Montelione, G. T.; Stuart, D. I.; Owens, R. J.; Daenke, S.; Schutz, A.; Heinemann, U.; Yokoyama, S.; Bussow, K.; Gunsalus, K. C., Protein production and purification. Nat Methods 2008, 5 (2), 135–46.

32. Savitzky, A.; Golay, M. J. E., Smoothing and differentiation of data by simplified least squares procedures. Anal Chem 1964, 36 (8), 1627–1639.

33. Mayer, M.; Meyer, B., Group epitope mapping by saturation transfer difference NMR to identify segments of a ligand in direct contact with a protein receptor. J Am Chem Soc 2001, 123 (25), 6108–17.

34. Haselhorst, T.; Lamerz, A. C.; Itzstein, M., Saturation transfer difference NMR spectroscopy as a technique to investigate protein-carbohydrate interactions in solution. Methods Mol Biol 2009, 534, 375–386.

35. Bhunia, A.; Bhattacharjya, S.; Chatterjee, S., Applications of saturation transfer difference NMR in biological systems. Drug Discovery Today 2012, 17 (9), 505–513.

36. Schrodinger, L., The PyMOL Molecular Graphics System. Version 2.0 Schrödinger, LLC.: 2015.

37. Laskowski, R. A.; MacArthur, M. W.; Moss, D. S.; Thornton, J. M., PROCHECK - a program to check the stereochemical quality of protein structures. J. App. Cryst. 1993, 26, 283–291.

38. Williamson, M. P., Using chemical shift perturbation to characterise ligand binding. Progress in Nuclear Magnetic Resonance Spectroscopy 2013, 73, 1–16.

39. Tornasz Cierpicki, J. O., Amide proton temperature coefficients as hydrogen bond indicators in proteins. Journal of Biomolecular NMR 2001, 21, 249–261.

40. Maier, J. A.; Martinez, C.; Kasavajhala, K.; Wickstrom, L.; Hauser, K. E.; Simmerling, C., ff14SB: Improving the Accuracy of Protein Side Chain and Backbone Parameters from ff99SB. J Chem Theory Comput 2015, 11 (8), 3696–713.

41. D.A. Case, I. Y. B.-S. S.R. Brozell, D.S. Cerutti, T.E. Cheatham, III, V.W.D. Cruzeiro, T.A. Darden, R. E. Duke, D. Ghoreishi, M.K. Gilson, H. Gohlke, A.W. Goetz, D. Greene, R Harris, N. Homeyer, S. Izadi, A. Kovalenko, T. Kurtzman, T.S. Lee, S. LeGrand, P. Li, C. Lin, J. Liu, T. Luchko, R. Luo, D.J. Mermelstein, K.M. Merz, Y. Miao, G. Monard, C. Nguyen, H. Nguyen, I. Omelyan, A. Onufriev, F. Pan, R. Qi, D.R. Roe, A. Roitberg, C. Sagui, S. Schott-Verdugo, J. Shen, C.L. Simmerling, J. Smith, R. Salomon-Ferrer, J. Swails, R.C. Walker, J. Wang, H. Wei, R.M. Wolf, X. Wu, L. Xiao, D.M. York and P.A. Kollman Amber 2018, University of California, San Francisco., 2018.

42. Feig, M.; Karanicolas, J.; Brooks, C. L., 3rd, MMTSB Tool Set: enhanced sampling and multiscale modeling methods for applications in structural biology. J Mol Graph Model 2004, 22 (5), 377–95.

43. Schrodinger, 9.0; LLC, New York, NY., 2009.

44. Friesner, R. A.; Banks, J. L.; Murphy, R. B.; Halgren, T. A.; Klicic, J. J.; Mainz, D. T.; Repasky, M. P.; Knoll, E. H.; Shelley, M.; Perry, J. K.; Shaw, D. E.; Francis, P.; Shenkin, P. S., Glide: a new approach for rapid, accurate docking and scoring. 1. Method and assessment of docking accuracy. J Med Chem 2004, 47 (7), 1739–49.

45. Humphrey, W.; Dalke, A.; Schulten, K., VMD: visual molecular dynamics. J Mol Graph 1996, 14 (1), 33-8, 27-8.

